# Ocular dominance columns in mouse visual cortex

**DOI:** 10.1101/2023.07.22.550034

**Authors:** Pieter M. Goltstein, David Laubender, Tobias Bonhoeffer, Mark Hübener

## Abstract

The columnar organization of response properties is a fundamental feature of the mammalian visual cortex. However, columns have not been observed universally across all mammalian species. Here, we report the discovery of ocular dominance columns in mouse visual cortex. Our observation in this minute cortical area sets a new boundary condition for models explaining the emergence of columnar organizations in the neocortex.

## Introduction

Cortical columns have traditionally been proposed to represent basic anatomical and functional units tessellating the mammalian neocortex^1, 2^. Within the vertically oriented columns, neurons across cortical layers share functional properties, while across the cortical surface these properties typically change gradually, with occasional abrupt jumps, forming maps consisting of repetitive modules^3, 4^. In the primary visual cortex (V1), this architecture gives rise to, for instance, the orientation preference map and ocular dominance columns^5, 6^. However, while several maps have been found in mammals such as primates^1, 7, 8^ and carnivorans (e.g. cats^9^ and ferrets^10^), in rodents it is less clear to what extent the visual cortex is functionally organized.

Interestingly, the first electrophysiological studies in mouse V1 did observe a certain degree of functional clustering of orientation^11^ and possibly also eye preference^12^. However, later work using two-photon calcium imaging did not find any obvious maps for these features^13–15^ and it was a widely held belief that maps in visual cortex were largely absent in the mouse. More recently, some functional clustering at the micro-scale^16, 17^, fluctuations in the density of ON/OFF neurons^18^, and potentially a global organization for orientation preference spanning multiple visual areas^19^ were reported in mouse visual cortex. In contrast, in rat binocular V1, a pattern of ipsilateral eye domains, most prominently identifiable in cortical layer 4, was reported using immediate early gene labeling and electrophysiology^20–22^. Thus, we asked whether a pattern of ocular dominance columns in mice had just been overlooked, or whether it in fact does not exist, potentially because mouse binocular V1 is just too small a cortical area to harbor such an elaborate functional architecture.

## Results

In order to map visual cortex function, we used low-magnification two-photon calcium imaging in GCaMP6s transgenic mice^23^ (n=9), recording neuronal activity over an area of approximately 1 mm^2^, covering mouse binocular as well as monocular V1 (Fig. 1a). Orientation tuning and ocular dominance of layer 4 neurons were assessed by presenting drifting gratings, moving in one of eight possible directions, to each eye independently (Fig. 1a,b). The resulting layer 4 volumetric recordings, spanning four imaging planes (spaced 20 μm apart), contained on average 5804 (± 1042 s.d.) visually responsive (stimulus-tuned; see Methods) neurons per mouse, with 4435 (± 761 s.d.) neurons preferring the contralateral eye and 1369 (± 517 s.d.) neurons preferring the ipsilateral eye. The ocular dominance index (ODI; ranging from -1 to +1, ipsi to contra; see Methods) of visually responsive neurons was skewed to the contra eye (Fig. 1c,d) having a mean of 0.31 (± 0.09 s.d) across animals.

**Figure 1:**
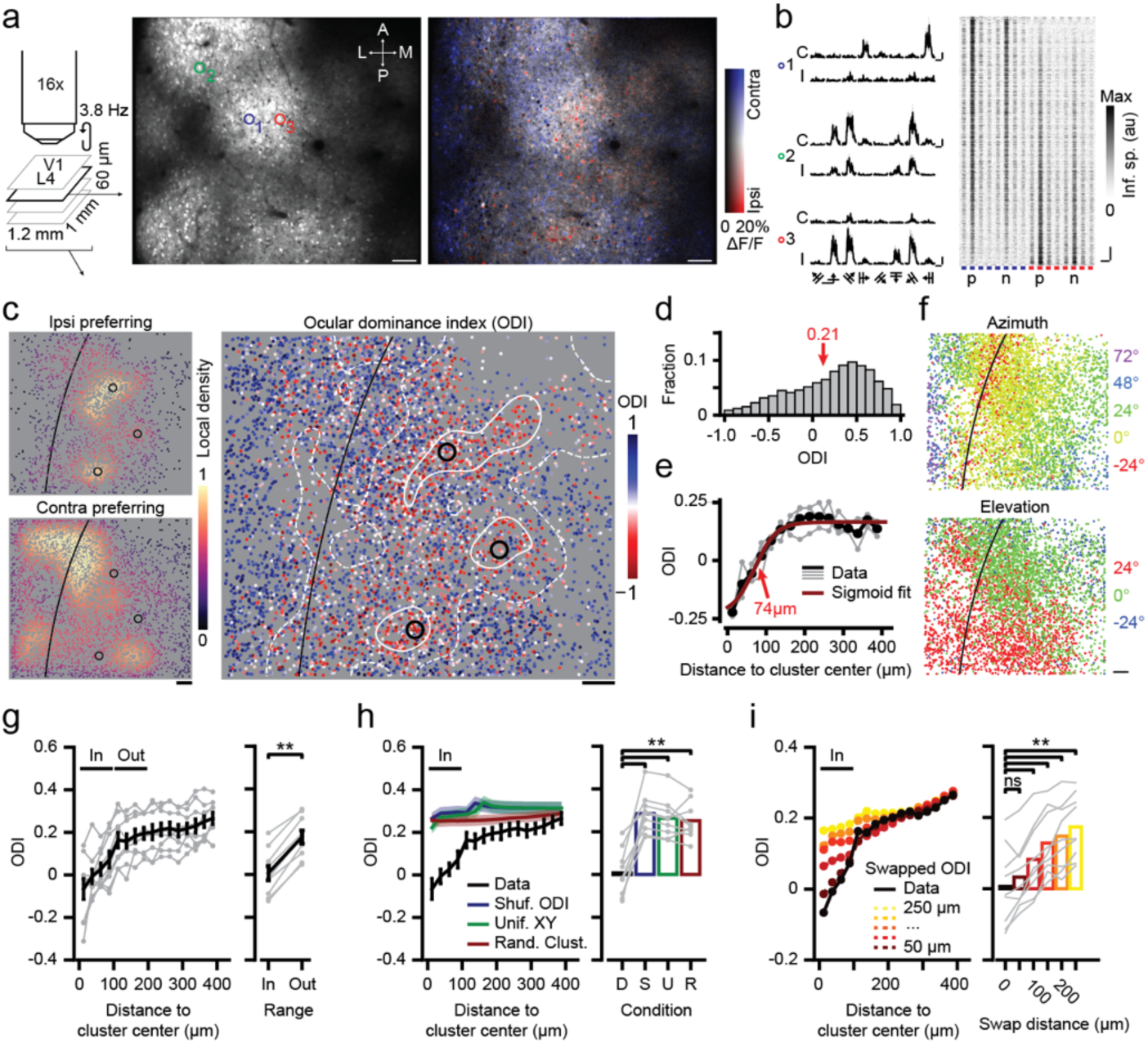
Spatial clustering of ocular dominance in layer 4 of mouse visual cortex. **a,** Multiplane, large-field of view (FOV) two-photon calcium imaging of ocular dominance in layer 4 of the visual cortex. Left: Schematic indicating volume dimensions and acquisition rate. Middle: Example FOV (GCaMP6s expression) of a single plane; ‘A’,’P’,’M’ and ’L’ indicate anterior, posterior, lateral and medial. Right: HLS map for ocular dominance. Hue: Eye-preference (contralateral: Blue; ipsilateral: Red); Lightness: ΔF/F; Saturation: Eye selectivity. Scale bar: 100 μm. **b,** Left: ΔF/F response to drifting grating stimuli for three example cells (see **a**). “C”: Contralateral, “I”: Ipsilateral. Scale bar, vertical: 100% ΔF/F, horizontal: 10 seconds. Right: Inferred spiking activity of all visually responsive cells (n=1834) in **a**, sorted vertically by ODI and aligned horizontally to the preferred (P) direction (N: Null direction; blue/red: Contra/ipsi). Scale bar, vertical: 100 neurons, horizontal: 10 seconds. **c,** Left: Local density of ipsilateral (top) and contralateral (bottom) eye, preferring visually responsive neurons separately (across four-plane volume). Black line: lateral boundary of V1 (V1 is to the right). Black circles: Ipsi-cluster centers (see Methods). Right: All visually responsive neurons, color coded for ODI. White equi-ODI lines indicate ODI=0 (solid) and ODI=0.2 (dashed). **d,** Histogram of ODI (for volume in **c**). **e,** Mean ODI as a function of distance to the three ipsi-cluster centers in **c** (individual ipsi- clusters in gray). Red line: sigmoid fit; the point of maximum inclination (red arrow) approximates the ipsi- cluster radius. **f,** Preferred azimuth (top) and elevation (bottom) of the neurons shown in **c**. Scale bar: 100 μm. **a-f**, Data of mouse M02. **g,** Left: ODI as function of distance to ipsi-cluster centers. Black: Mean ± s.e.m., gray: Individual mice (n=9). Right: Mean ODI inside (“In”, 0-100 μm) and outside (“Out”, 100-200 μm) ipsi-clusters (two-sided WMPSR test, W=0, p=0.004, n=9 mice). **h,** Same as **g**, black line (mean ± s.e.m.) shows actual data (‘D’), blue, green and red lines show global randomization controls. Right: Mean (± s.e.m.) ODI “In” ipsi- clusters for real and shuffled data (two-sided Kruskal-Wallis test, H3=17.7, p=5.0·10^-04^, post hoc two-sided WMPSR test, **p<0.01, n=9 mice). Blue, ‘S’: ODI values shuffled across neurons. Green, ‘U’: XY coordinates of neurons resampled from uniform distribution. Red, ‘R’: Ipsi-cluster centers randomly placed in FOV. **i,** As **h**, colored lines show local randomization controls in which the positions of pairs of neurons, spaced at a distance of 50 μm (dark red) to 250 μm (yellow), were swapped (two-sided Kruskal-Wallis test, H5=16.8, p=0.0048, post hoc two-sided WMPSR test, ns: not significant, ** p<0.01, n=9 mice).

### Clusters of ipsilateral eye preferring cells in cortical layer 4

When inspecting HLS (hue, lightness, saturation) maps for ocular dominance (Fig. 1a), we found that some mice showed clear patches of cells with ipsilateral eye preference (Fig. 1c). In order to quantify these clusters, we used a local-density based clustering algorithm^24^, allowing to identify patches of cells responding preferentially to the ipsilateral eye (ipsi-clusters; see Methods). We calculated the average ODI across all neurons (both ipsi- and contra-preferring) as a function of distance to the centers of the detected ipsi-clusters. If ipsi-clusters merely reflected an overall uneven spatial distribution of neurons, there should be a similar pattern for contra preferring cells, and the average ODI near ipsi-clusters should be relatively similar to its surround. However, the mean ODI near ipsi- clusters was just below zero (-0.07 ± 0.15 s.d.), indicating that most nearby neurons indeed responded preferentially to the ipsilateral eye. Furthermore, ODI increased with distance from the cluster center, showing that neurons outside the ipsi-clusters generally preferred the contralateral eye (Fig. 1e, g).

Comparison with single neuron-resolution retinotopic maps revealed that the ipsi-clusters were located within binocular V1 (Fig. 1f; Figs. S1, S2). Individual ipsi-clusters varied in shape from being round to elongated, and occasionally had irregular features. All ipsi-clusters (average size: 162 μm, ± 26 μm s.d.; see Methods) were substantially smaller than the extent of binocular V1, and in most mice we detected multiple ipsi-clusters (2.7 ± 0.7 s.d ipsi-clusters per mouse) within the approximately 50% of binocular V1 that our field of view covered on average.

We performed several global randomization procedures to test whether ipsi-clusters could occur by chance: neither when shuffling the ODI values across neurons, when randomly repositioning the ipsi-cluster centers, nor when repositioning neurons at random XY coordinates in the imaged region did we observe ipsi-clusters having ODI values comparable to the real data (Fig. 1h). This shows that ipsi-clusters, as observed, do not emerge from random spatial distributions of ipsilateral eye preferring cells.

Because our imaging field of view (FOV) was wider (∼1.2 mm) than the medial-lateral extent of the binocular visual cortex (∼0.8 mm), it could be that the effect in Fig. 1h, at least partly, reflected the monocularity of neurons outside the binocular visual cortex. To address this issue, we tested whether the ipsi-clusters were part of a fine-grained spatial organization for ocular dominance, smaller than the extent of binocular V1, or whether they could be explained by global effects like boundaries between binocular and monocular cortex. This was done by a local randomization control. By swapping the ODI values of pairs of neurons that were separated by a small distance (e.g. 50 μm), we randomized the local spatial distribution of ODI values while maintaining the larger-scale functional organization of monocular and binocular visual cortex (Fig. S3). This analysis showed that the ipsi-clusters indeed originated in the fine-grained arrangement of ipsilateral eye preferring neurons (Fig. 1i).

In order to exclude that the functional clustering of ocular dominance described here is a specific feature of the transgenic mouse line used, we confirmed our findings in a different line using virus-mediated (AAV2/1) expression of a red-shifted calcium indicator (jRGECO1a) in the right hemisphere of Scnn1a-Tg3-Cre transgenic mice, thus limiting expression in V1 to cortical layer 4 (see Methods). Despite small quantitative differences, which likely resulted from a less homogeneous distribution and overall smaller number of neurons expressing the calcium indicator, we confirmed the overall finding of ipsi-clusters in binocular V1 (Fig. S4).

### L4 ipsi-clusters extend vertically into cortical layer 2/3 and 5

Having found clusters of ipsilateral eye preferring neurons in layer 4 of mouse binocular V1, we asked whether these ipsi-clusters extended vertically into other cortical layers. In each animal (n=9), we acquired an imaging volume spanning 360 μm of cortex from upper L2/3 (170 μm below the pial surface) to upper L5 (530 μm; 37 planes, spaced 10 μm apart; see Fig. 2a). The volume was constructed from 12 individual four-plane imaging stacks, which were acquired in random order (see Methods). Each L2/3-L5 volume contained on average 24366 (± 6384 s.d.) visually responsive neurons, with 18305 (± 5879 s.d.) neurons preferring the contralateral eye and 6061 (± 2600 s.d.) preferring the ipsilateral eye and a mean ODI of 0.27 (± 0.11 s.d.) across all imaged cortical layers.

**Figure 2:**
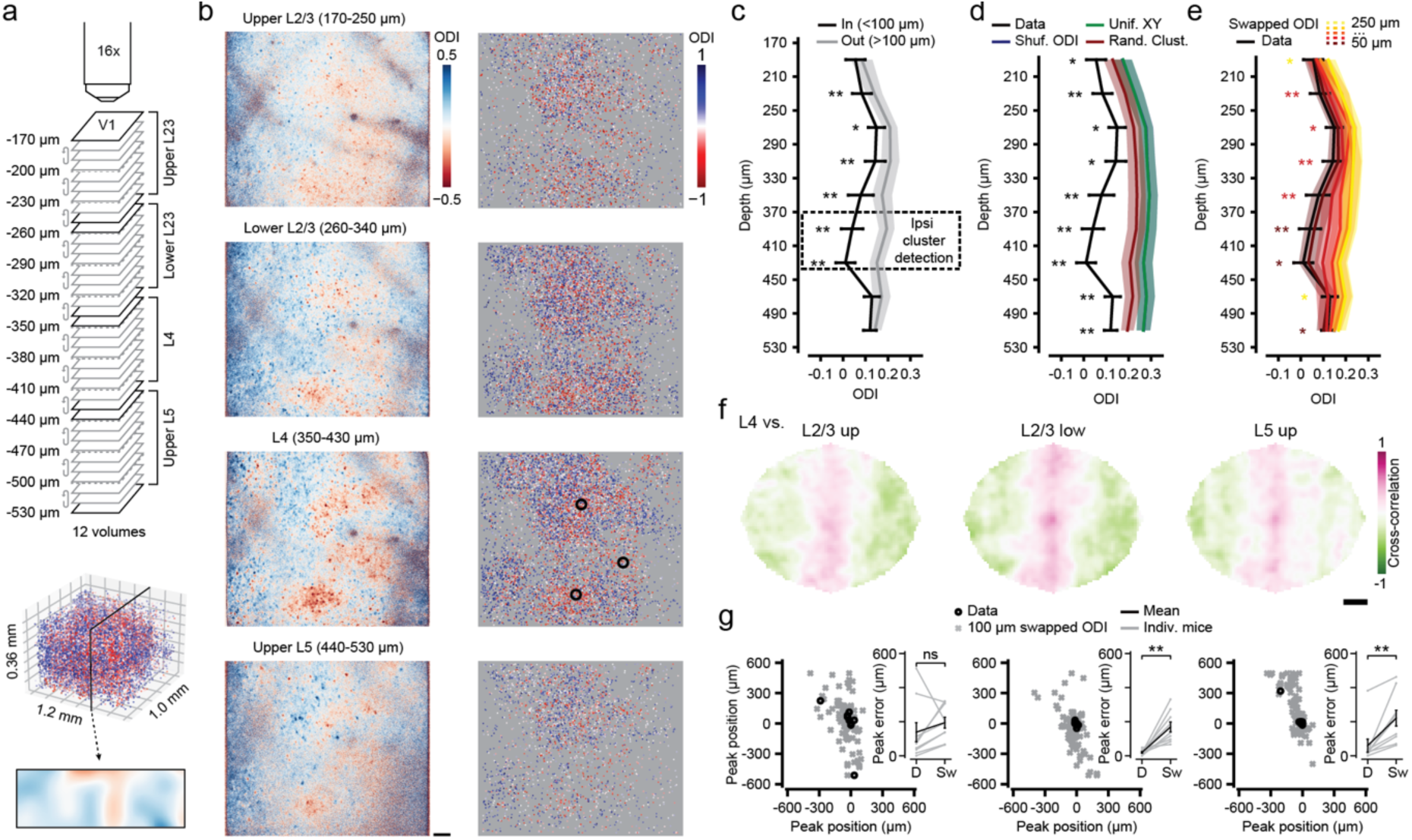
Ocular dominance columns in mouse visual cortex layers 2/3, 4 and 5. **a,** Approach for recording ocular dominance across a cortical volume. Top: Schematic showing 12 multilevel imaging stacks (acquired in random order) resulting in 37 uniquely imaged planes, spaced 10 μm apart in depth. Middle: ODI of all visually responsive neurons in an example volume (n=22898; color bar: see **b**, right). Bottom: Projection of ODI across a 100 μm thick vertical slice (color bar: see **b**, left). **b,** Left: Pixelwise ODI maps using imaging data combined across multiple imaging planes spanning four depth ranges (see **a**). Scale bar: 100 μm. Right: ODI of all visually responsive neurons across the same depth ranges. Black circles: Ipsi- clusters. **a,b,** Data of mouse M02. **c,** Mean (± s.e.m. across 9 mice) ODI “In” (<100 μm; black) and “Out” (100 μm-200 μm; gray) of ipsi-clusters, detected in layer 4 (dashed box), across nine depth bins (tick marks indicate bin-edges). **d, e,** As **c**, real data (mean ± s.e.m., black) “In” ipsi-clusters versus global and local randomization controls. **d**, Global randomization control. Blue: Shuffled ODI. Green: Uniformly resampled XY coordinates. Red: Randomly placed ipsi-cluster centers. **e**, Local randomization control. Dark red to yellow mark swap distances from 50 μm to 250 μm. **c-e,** Statistical comparison of ODI “In” ipsi-cluster centers versus all controls (“Out”, local and global shuffles), Kruskal-Wallis tests per depth bin, p<0.05, corrected for 9 comparisons; post hoc two-sided WMPSR tests, *p<0.05, **p<0.01, in **e** color coded for smallest significant swap distance, n=9 mice). **f,** 2D cross-correlation of L4 ODI maps with those of L2/3 and L5 (see Methods; see Fig. S7a; Data of mouse M02). **g,** Spatial position of the cross-correlation peak (real data, ‘D’, black) compared to locally randomized data (‘Sw’, swapped ODIs at 100 μm, 10 repeats, gray). Left: Upper L2/3, middle: Lower L2/3, right: Upper L5. Inset shows the peak error, i.e. the Euclidian distance between the detected peak and the center of the cross-correlation map (L2/3 up: Two-sided WMPSR test, W=10, p=0.16; L2/3 low: Two-sided WMPSR test, W=0, p=0.004; L5 up: Two-sided WMPSR test, W=0, p=0.004; n=9 mice; ns: not significant, ** p<0.01).

In several mice, pixelwise ODI maps across different cortical depths showed a clear similarity in the overall patterns of ipsilateral and contralateral eye dominated regions (Fig. 2b, left; Fig. S5).

Using the method described above, we identified centers of ipsi-clusters in the L4 subvolume spanning 350 μm to 430 μm below the pial surface (Fig. 2b, right; see Methods). The ODI of neurons in a small column (<100 μm range around ipsi-cluster centers, <200 μm diameter) above and below the L4 ipsi-cluster centers was significantly lower (more ipsilateral) than the ODI of neurons further outwards (range: 100-200 μm, diameter: 200-400 μm; Fig. 2c; Fig. S6). As within layer 4, the global and local shuffle controls showed that the low ODI in the column above and below the L4 ipsi-cluster centers did not occur by chance, nor did it reflect the division of visual cortex in monocular and binocular regions (Fig. 2d,e; Fig. S6). Thus, the ipsi-clusters we detected in cortical layer 4 extended vertically, in a columnar fashion, at least into cortical layers 2/3 and 5.

In order to directly visualize how the spatial organization of ODI in layer 4 extended across cortical layers, we constructed high-resolution ODI-maps based on cellular ODI values for four depth ranges (corresponding to upper and lower L2/3, L4 and upper L5; see Methods; Fig. S7a). The similarity in ODI patterns between L4 and other layers was calculated as the two-dimensional cross-correlation between ODI-maps (Fig. 2f; Fig. S7a,b). The peaks of nearly all cross-correlation maps were close to the origin (0,0), confirming the alignment of the overall ODI pattern across layers (Fig. 2g, black circles; Fig. S7c,d). In comparison, the locations of cross-correlation peaks for ODI maps calculated using globally shuffled ODI values (shuffled ODI), and using imaging planes from altogether different animals (shuffled planes) were randomly distributed (Fig. S7c,d). The cross-correlation peaks for ODI maps calculated using locally shuffled ODI values (ODIs swapped at 100 or 200 μm) were still positioned close to the center on the x-axis, likely reflecting the overall binocular-monocular gradient along this dimension. In contrast, along the vertical image axis (y) there was much less of a coarse gradient in ODI values, and local randomization prevented the cross-correlation peaks to position near the center (Fig. S7c-e). This indicates that the alignment of ODI patterns across cortical layers depends on the fine-grained functional organization of cellular ocular dominance. We conclude that mouse visual cortex contains ocular dominance columns.

## Discussion

Finding columnar structures for ocular dominance in the minute mouse binocular visual cortex provides a new opportunity for investigating the general question of which factors determine whether a cortex has columns or not. Experimental^13, 25, 26^ and theoretical^27^ studies have argued that columnar architectures in the visual cortex are only found in certain mammalian orders, such as primates, carnivorans, ungulates, scandentians and diprotodonts, but not in others. Rodent visual cortex, in particular, has been thought to lack columnar organizations. Our and others’ recent findings of ocular dominance (rat^20–22^) and ON/OFF (mice^18^) domains show that this distinction based on taxonomy does not hold. For another prominent columnar system in the visual cortex, the orientation preference map, the situation is less clear. So far, no such map has been found in any rodent. However, preliminary data indicate a large-scale organization for preferred orientation across all of mouse visual cortex^19^, and orientation-minicolumns have been found in several rodent species^16, 17, 26, 28^. Importantly, orientation and direction preference maps have been observed in another part of the mouse visual pathway, the superior colliculus^29–32^ (see however^33^). Thus, circuits in the visual system of the mouse and rat are in principle capable of organizing into columnar structures. Studies in a larger variety of rodent species might reveal whether their visual cortex can also hold a “proper” orientation map, like those found in cats and primates^7, 9^.

Apart from taxonomy, factors like visual cortex size^34^, the absolute number of neurons^35^ or the retino-thalamo-cortical mapping ratio^36–38^ have been put forward in theoretical work to explain the absence of columns in a visual cortex as small as that of the mouse. While some of these studies refer to a columnar architecture in general, or orientation columns specifically, others^35^ explicitly predict that mouse visual cortex does not have ocular dominance columns, contrary to what our experiments show. Our finding will help improving future models of visual cortex functional architecture.

In cats and monkeys, during early development, ocular dominance columns have been shown to gradually emerge from initially intermingled thalamic axons driven by one or the other eye^39, 40^. This process is generally thought to be governed by neuronal activity, either visually driven or internally generated^41^, and serves as a prime example for activity dependent development in the nervous system. There are, however, alternative (or additional) explanations, which point to molecular axon guidance cues being an important factor for the formation of ocular dominance columns^42, 43^. Research on this topic has not progressed much recently, since the experimental work was largely performed in ferrets and cats. These species do not lend themselves easily to genetic interventions, which may be crucial to elucidate the molecular nature of such putative guidance cues. The fact that we have now demonstrated ocular dominance columns in the mouse visual cortex makes such experiments feasible and very worthwhile.

Our finding expands the range of mammalian species, across very diverse orders, which display ocular dominance columns. That ocular dominance columns are apparently rather the rule than an exception is in stark contrast to the lack of a clear hypothesis of what their function for visual processing is, if there is one at all^44^. Possible explanations range from intracortical wirelength minimization^27, 35, 45^, over merely being an epi-phenomenon created by the activity dependent wiring of cortical circuits^46^, to a not very clearly spelled out function for binocular integration and stereoscopic depth perception^47^. A general way to probe the function of a structure in the brain is to remove it, and test for ensuing changes in neuronal processing and behavior. This appears very difficult for ocular dominance columns, since eliminating the columnar architecture without massively affecting cortical circuitry altogether seems impossible. There is, however, an experiment by nature, which might prove helpful for answering this question. Squirrel monkeys show a “capricious” expression of ocular dominance columns in their visual cortex, ranging from fully developed columns in some animals to nearly complete absence in others^48^. While we have not systematically explored the variability in the degree of columnar organization in our mouse data, there are clear differences between individual animals. Recent experiments have shown that mouse visual cortex contains many neurons sensitive to binocular disparities^49, 50^, and that mice make use of such binocular cues for judging distances^51–53^.

Relating the degree of columnar organization for ocular dominance to neuronal or behavioral signs of binocular depth perception might reveal whether the arrangement of cortical neurons into eye specific columns is relevant for these important visual system functions.

## Methods

### Animals

All experiments were conducted following the institutional guidelines of the Max Planck Society and the regulations of the local government ethical committee (Beratende Ethikkommission nach §15 Tierschutzgesetz, Regierung von Oberbayern). Nine adult mice (6 female, 3 male; four to five months of age during data acquisition) genetically expressing the calcium indicator GCaMP6s in excitatory neurons (B6;DBA-Tg(tetO-GCaMP6s) 2Niell/J^23^, JAX stock #024742; back-crossed for at least seven generations to C57Bl/6NRj) crossed with B6.Cg-Tg(Camk2a-tTA)1Mmay/DboJ^54^ (JAX stock #007004; maintained on a mixed background of C57BL/6NRj and C57BL/6J) and seven adult Scnn1a-Tg3-Cre mice (4 female, 3 male; four months of age during data acquisition; B6;C3- Tg(Scnn1a-cre)3Aibs/J^55^, JAX stock #009613; kept on a mixed background of C57BL/6 and C3H) were housed in small groups, or individually in case of inter-male aggression, in large cages (GR900, Tecniplast) containing a running wheel, a tunnel and a shelter. The animals were kept on a reversed day/night cycle with lights on at 22:00 h and lights off at 10:00 h. Food and water were available *ad libitum*.

### Surgery

Animals were anesthetized with a mixture of fentanyl (0.05 mg/kg), midazolam (5.0 mg/kg) and medetomidine (0.5 mg/kg) in saline (injected i.p.; FMM in short). For analgesia, carprofen (5.0 mg/kg) was injected s.c. and lidocaine (0.2 mg/ml) was applied topically. A head bar and a 4 mm diameter cranial window (cover glass, #1 thickness) were implanted as described before^56, 57^. In Scnn1a-Tg3-Cre mice, AAV2/1.Syn.Flex.NES-jRGECO1a.WPRE.SV40 (titer: 2.6·10^13^; a gift from Douglas Kim & GENIE Project, Addgene viral prep #100854-AAV1) was pressure-injected using a glass micropipette at ∼400 μm depth (200–250 nl per injection), at 4 to 6 locations spanning binocular V1 (identified using intrinsic optical signal imaging)^58^. Following surgery, animals received antagonists (1.2 mg/kg naloxone, 0.5 mg/kg flumazenil and 2.5mg/kg atipamezole in saline, injected s.c.). Post-operative analgesia (5.0 mg/kg carprofen injected s.c.) was given for two subsequent days. In a subset of mice, small patches of bone-growth under the window were removed in a second surgery.

### Imaging

*In vivo* calcium imaging was performed using a customized, commercially available Bergamo II (Thorlabs, Germany) two-photon laser scanning microscope^59^ with a pulsed femtosecond Ti:Sapphire laser (Mai Tai HP Deep See, Spectra physics), running Scanimage 4^60^. GCaMP6s^61^ was excited using a wavelength of 940 nm, and jRGECO1a^62^ using 1050 nm. Lowpass (720/25 nm; Semrock, USA) and bandpass (GCaMP6s: 525/50-25 nm; jRGECO1a: 607/70-25 nm; Semrock, USA) filtered emitted fluorescence was detected with two GaAsP detectors (Hamamatsu, Japan). Laser power under the objective ranged from 15 to 45 mW, depending on the depth of imaging (upper L2/3 to L5). The field of view size of a single-plane image was 1192 × 1019 μm (XY; 1024 × 1024 pixels). Four-plane volumes were acquired at 3.8 Hz per plane using a 16x objective (0.8 NA; Nikon) attached to a piezo electric stepper (Physik Instrumente, Germany).

During imaging, animals were lightly anesthetized with FMM (see above) and kept warm on a heat pad (closed loop temperature controller set to 37°C). Eyes were kept moist using eye drops (Oculotect). Visual stimuli were presented on a gamma corrected and curvature corrected^63^ computer monitor. Ocular dominance and orientation tuning were assessed by presenting full screen, 100% contrast, square wave drifting gratings (spatial frequency: 0.04 cycles/degree; temporal frequency: 1.5 cycles/second) moving in one of eight possible directions and presented to the ipsilateral and contralateral eye separately using motorized eye-shutters. Stimulus presentation lasted 5 s, followed by a 6 s intertrial interval (ITI). Eye shutter switches were done 6 s post stimulus offset, and were followed by a 10 s post-switch interval, after which the next ITI started. Trials were presented in blocks of 16 stimuli, containing two blocks of eight movement directions, that is, one block for each eye. The eight directions were randomized per trial block, as the order of eye blocks. Each full block of 16 stimuli was presented 10 times in L4 experiments and five times in L2/3-L5 experiments.

For mapping retinotopy, drifting gratings having one of four (cardinal) directions were presented in subsections of the visual field (patches) measuring 26 × 26 retinal degrees (azimuth × elevation). The patches were centered on -48, -24, 0, 24 and 48 degrees azimuth and -24, 0 and 24 degrees elevation. Individual trials were grouped into blocks containing all combinations of 15 patches (visual field partitions) and 4 movement directions (thus 60 unique stimuli) in randomized order. Four blocks were presented per experiment, with each stimulus presentation lasting 4 s and each ITI lasting 5 s.

### Response maps

To visualize cortical response properties throughout a single field of view, we calculated pixel-wise ODI (ocular dominance index) and HLS (Hue, Lightness, Saturation) maps (see Fig. 1b and 2a for examples). First, stimulus fluorescence (F) images were produced by averaging images across all trials of each stimulus (from stimulus onset to stimulus offset) and a baseline F image was created by similarly averaging images acquired during the intertrial interval periods (from 0.9 ITI length before stimulus onset until stimulus onset). The stimulus and baseline F images were smoothed using a median filter (disk kernel, 1 pixel radius). Next, ΔF/F response maps were created for each stimulus by subtracting the baseline F image from the stimulus F image and dividing the result by the baseline F image. Negative and infinite values (caused by division by zero) were set to zero. HLS maps for ocular dominance were created by assigning for each pixel: (1) The color, hue (H), reflecting which eye resulted on average in the largest ΔF/F response (red for ipsilateral and blue for contralateral), (2) the brightness, lightness (L), reflecting the amplitude of the largest ΔF/F response, and (3) the saturation (S), reflecting the selectivity for either eye (similar to the ODI). ODI maps were created similarly, but here only the ODI value (see Eq. 1 below) per pixel was encoded according to a colormap (see legend next to maps).

### Image processing and source extraction

Volumetric imaging stacks were analyzed per plane using customized scripts (see Code availability) utilizing Suite2p version 0.7^64–66^. Image preprocessing consisted of dark-frame subtraction, line shift correction and image registration (both rigid and non-rigid). Source extraction and spike inference was done using the Suite2p algorithm (with ‘tau’ set to 1.5 s) and all further analyses were performed on the resulting inferred spiking activity per neuron. For L2/3-L5 data sets, the last and first plane of each pair of consecutive stacks were imaged at the same cortical depth, but data of only one of those planes (last) were used for analysis. Of neurons that occurred at the same XY position (centroid within a 6 μm diameter circle around another centroid) in consecutive planes, only the most significant tuned neuron (lowest p-value for stimulus tuning, see below) was kept.

### Tuning curve analysis

For each neuron and each trial, an average inferred spike response was calculated by subtracting the average inferred spike activity during baseline (1.8 s before stimulus presentation until stimulus onset; seven imaging frames) from the average inferred spiking activity during stimulus presentation. The trial-wise average inferred spike responses were grouped by stimulus features (eight directions × two eyes, or five azimuths × three elevations). We determined whether a neuron was significantly tuned to one or more stimulus features using a Kruskal-Wallis test, testing for differences in average inferred spike responses across stimuli (the alpha was set to 0.05). Throughout the manuscript, such stimulus-tuned neurons are referred to as “visually responsive neurons”. For each visually responsive neuron recorded in ocular dominance and orientation tuning sessions, we quantified the ocular dominance index (ODI; Eq. 1) using the average inferred spiking response (*R*) to the preferred direction (*pref dir*) of each eye (*ipsi* or *contra*).

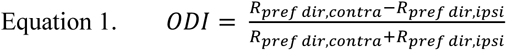

In addition, we calculated the preferred orientation and direction from the stimulus that elicited the largest inferred spiking response (thus based on the response to the preferred eye). For neurons in retinotopic mapping experiments, we calculated the preferred azimuth and elevation using the same approach as for orientation and direction. Finally, all significantly tuned neurons were grouped in a single volume spanning 60 μm in layer 4 or 360 μm from upper layer 2/3 to layer 5.

### Ipsi-cluster detection

Volumes were split in a set of ipsilateral and contralateral eye preferring neurons by thresholding at an ODI value of 0. For layer 4 volumes, all significantly visually responsive neurons were included, and for layer 2/3 to layer 5 volumes only neurons between 370 and 430 μm in depth were included. Clusters of ipsilateral eye preferring neurons were identified using the ‘fast search and find of density peaks’ algorithm^24^. In brief, the local density for neuron *i* (ρ*_i_*) was calculated following Eq. 2, where *N* equals the total number of neurons, *d_ij_* is the Euclidian distance of neurons *i* and *j* (μm in cortical space), and *d_c_* is a cutoff-distance set to the fifth percentile of the distribution of all local distances.

The function *χ*(*x*) returns 1 when *x* assumes values below zero, otherwise it returns 0.

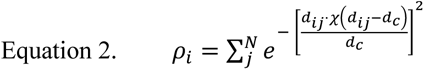

Next, the distance of each neuron *i* to the nearest point of higher local density (δ*_i_*) was determined following Eq. 3.

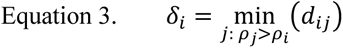

Peaks in the local density of ipsilateral eye preferring neurons, ipsi-cluster centers, were identified as neurons with an ρ*_i_* exceeding 0.2 · ρ*_max_*, and a δ*_i_* exceeding 0.2 · δ*_max_*. Typically, one to five ipsi-cluster centers were detected in each L4 volume, which spanned roughly 50 percent of the binocular visual cortex.

### Randomization procedures

We performed several randomization controls. For the “Shuffled ODI” control we randomized ODI values with respect to neuron identity. For the “uniform XY” control we drew for each neuron new XY coordinates from a uniform distribution ranging from the minimum to maximum of the original set of neuron positions. For the “Random clusters” control we drew new XY coordinates for the originally detected ipsi-clusters, also from a uniform distribution, but with the range set 200 μm inwards from the minimum and maximum of the neuron positions. Finally, for the “Swapped ODI” control we iteratively selected the pair of neurons, without replacement, that was spatially separated by a distance closest to a specified reference distance (50 μm, 100 μm, .., 250 μm) and swapped their ODI values. We continued this process until all neurons were paired up and had their ODI values swapped (typically five to 20 final pairs were separated by distances deviating more than 50 μm from the specified distance). In all controls, ipsi-cluster centers were identified anew after the randomization procedure.

#### Cross correlation maps

Spatial auto- and cross-correlation maps were calculated based on smoothed single-neuron ODI maps, which were constructed as follows. A ‘summed ODI map’ was calculated by, for each pixel, summing the ODI values of all neurons that had their centroid at that pixel. Next, a ‘coverage map’ was constructed, in which the value of each pixel reflected the number of neurons having their centroid at that pixel. Finally, a smoothed single-neuron based ODI map was obtained by dividing the ‘summed ODI map’ by the ‘coverage map’, and smoothing the result with a two-dimensional Gaussian kernel (σ = 16 μm). Using the pixel-wise Pearson product-moment correlation between two smoothed ODI maps at varying spatial offsets (Eq. 4) we obtained the auto and cross-correlation maps^43, 67^.

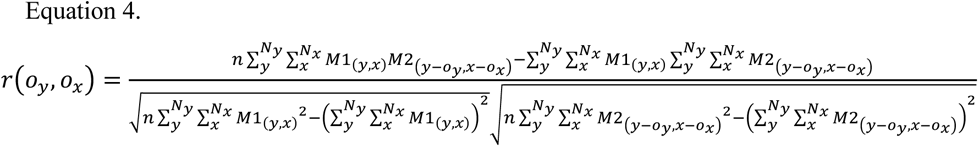

The cross-correlation *r* was calculated at each pixel offset (ο*_x_* and ο*_y_*) individually. *N_y_* and *N_x_* represent the number of pixels along the y and x dimensions in the smoothed ODI maps *M1* and *M2*, *n* is the total number of pixels in the map, and *M1_(y,x)_* returns the ODI value of map *M1* at pixel position (*y*,*x)*. The cross-correlation was only calculated for pixel-offsets at which minimally 20% of the smoothed ODI maps overlapped.

### Statistics

Data analysis and statistical testing was performed using Python (3.8.12), Numpy (1.20.3) and Scipy (1.7.3). We did not use statistical methods to predetermine the sample size, but used an *n* similar to that reported in previous publications^14, 15^. We excluded data of one Scnn1a-Tg3-Cre transgenic mouse (dataset shown in ED Fig. 4) because the signal-to-noise ratio was low and Suite2P only detected ∼300 neurons in the entire L4 volume, which was not sufficient for identifying densities of ipsilateral eye preferring neurons. Experimenters were not blinded to experimental conditions. All data are represented as mean (± s.e.m.) unless indicated otherwise. Group-wise differences were tested using non-parametric statistical tests with alpha set to 0.05.

## Acknowledgements

We thank Max Sperling, Volker Staiger, Claudia Huber, Frank Voss and Ursula Weber for technical assistance. This project was funded by the Max Planck Society and the Collaborative Research Center SFB870 (project number A08, reference number 118803580) of the German Research Foundation (DFG) to M.H.

## Author contributions

P.M.G. and M.H. designed the experiments. P.M.G. and D.L. conducted experiments. P.M.G. programmed analysis code and analyzed data. P.M.G., D.L., T.B. and M.H. discussed the data and wrote the manuscript.

## Data availability

Data supporting this study will be available on https://gin.g-node.org/pgoltstein/mouse-od-columns/.

## Code availability

The Python code used for data analysis and production of figures will be available on https://github.com/pgoltstein/mouse-od-columns/. Custom-written MATLAB and Python routines used for data collection and data preprocessing are available upon reasonable request.

---

**Figure S1:**
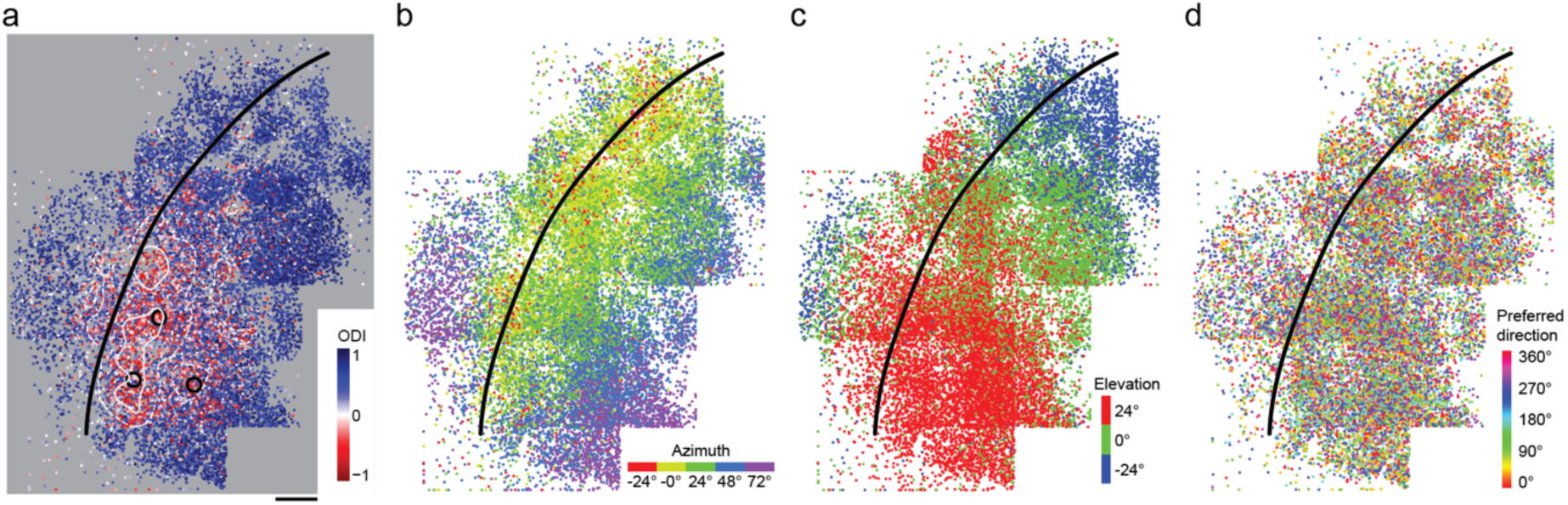
Tiled, large-field of view mapping of ocular dominance, retinotopy and orientation preference in cortical layer 4 of GCaMP6s transgenic mice. **a-d,** Spatial position of all visually responsive neurons from five multiplane layer 4 recordings, stitched into a single map color-coded for **a** eye preference, **b** preferred azimuth, **c** preferred elevation and **d** preferred orientation (data of mouse M06). Black line: Estimated boundary separating primary visual cortex (right) and lateral higher visual areas (left). White lines in **a** indicate ODI=0 (solid) and ODI=0.2 (dashed), black circles mark identified ipsi-clusters. Scale bar: 200 μm.

**Figure S2:**
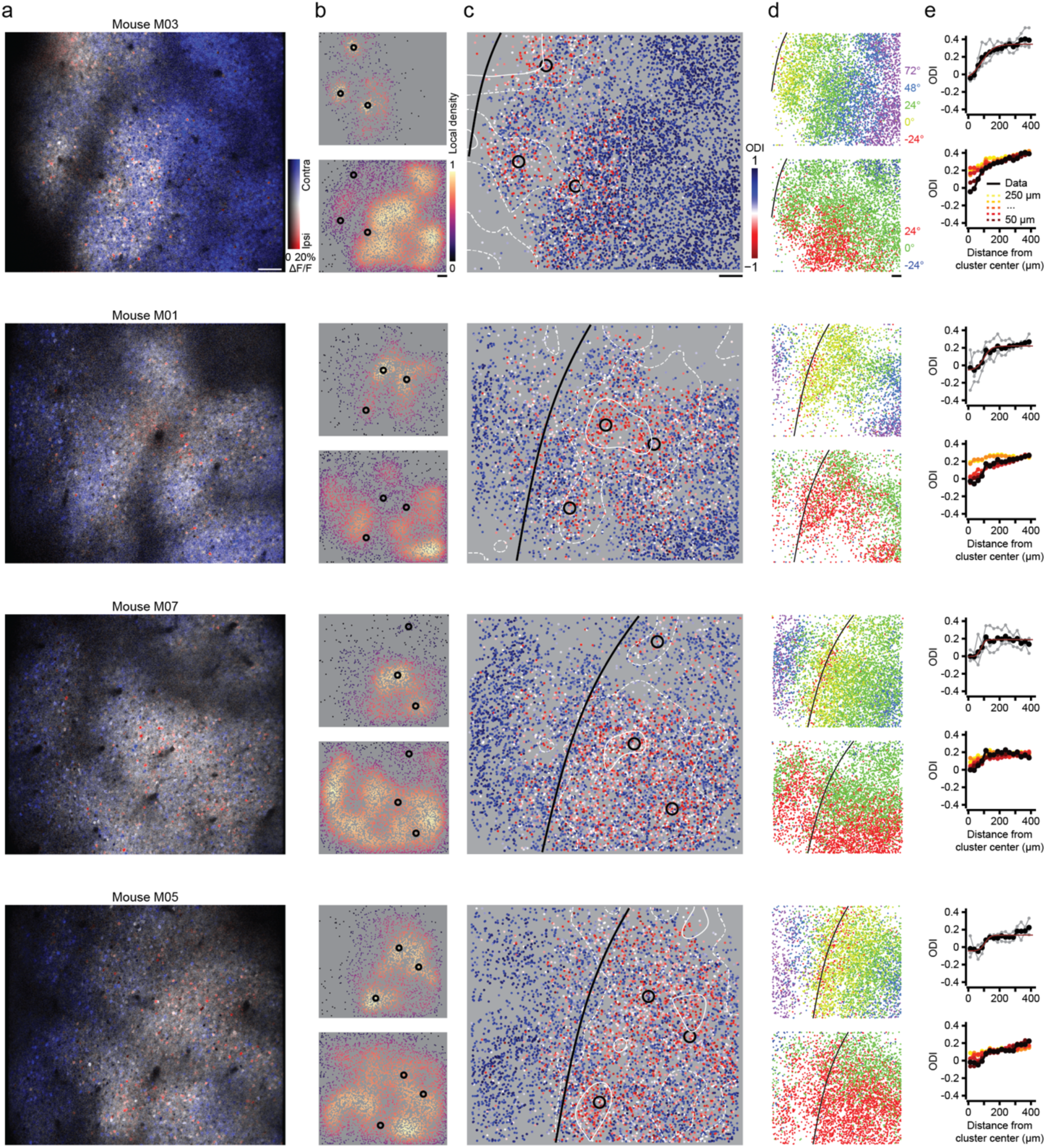
Examples of ipsilateral eye preferring ocular dominance clusters in GCaMP6s transgenic mice. Each row shows data of all visually responsive neurons recorded from four imaging planes in layer 4 of a single mouse (see Fig. 1a). **a,** HLS map for ocular dominance. Hue: Eye-preference (contralateral: Blue; ipsilateral: Red). Lightness: ΔF/F response amplitude. Saturation: Eye selectivity). **b,** Local density for each visually responsive neuron. Top: Ipsilateral eye preferring neurons. Bottom: Contralateral eye preferring neurons. Black circles mark the centers of ipsi-clusters, detected in the local density map for ipsilateral eye preferring neurons (top). **c,** Ocular dominance of each visually responsive neuron. White iso-ODI lines delineate ODI=0 (solid) and ODI=0.2 (dashed), the black line shows the lateral border of V1 (see **d**). **d,** Preferred azimuth (top) and elevation (bottom) in the layer 4 volume. Black line: V1 lateral boundary. **a-d,** Scale bar: 200 μm. **e,** Top: ODI as a function of distance to ipsi-cluster centers (Gray: Individual ipsi-clusters. Black: Mean across ipsi-clusters). Bottom: As top, but for the local randomization control, swapping the ODIs of pairs of cells at a radius of 50 μm (dark red) to 250 μm (yellow).

**Figure S3:**
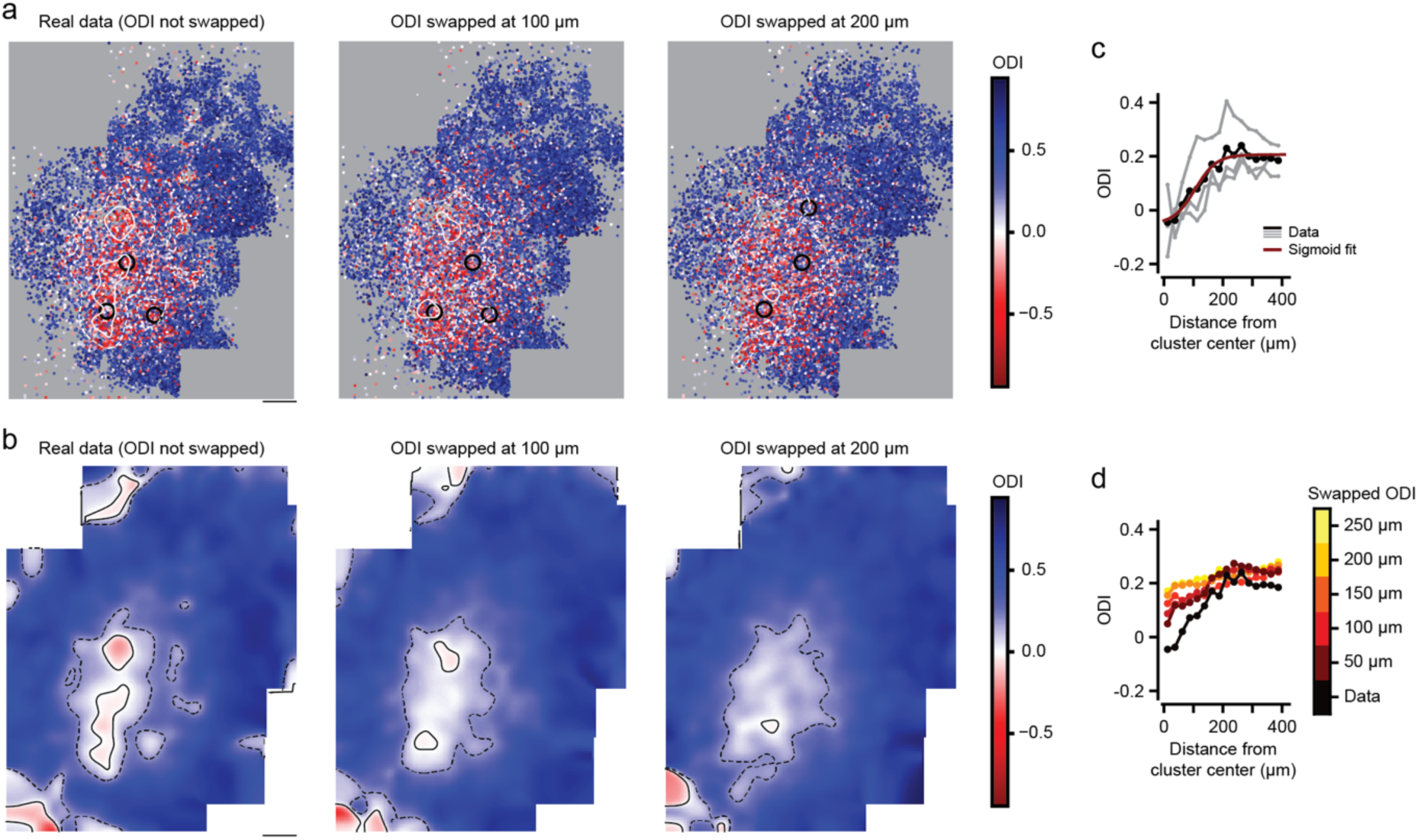
Local randomization by repositioning cells within a fixed radius of their original position. **a,** ODI of each visually responsive neuron in a tiled (five tiles), multiplane (four levels) L4 volume, stitched together to represent a large part of visual cortex including the complete binocular region and several higher visual cortical areas (see also Fig. S1). Left: Original data. Middle: Local randomization by swapping the ODIs of pairs of neurons at a distance of 100 μm. Right: As middle, but for swapping pairs of neurons at 200 μm distance. White: Iso-ODI lines for ODI=0 (solid) and ODI=0.2 (dashed). Black circles: Centers of ipsi-clusters detected in each data set. Data of mouse M06. **b,** Smoothed ODI maps, based on the data shown in **a**. Smoothing was done using a 50 μm Gaussian kernel. Black: Iso-ODI lines for ODI=0 (solid) and ODI=0.2 (dashed). **a,b,** Scale bar: 200 μm. **c,** ODI as a function of distance to ipsi-cluster centers for the data in **a**, left. Gray: Data for individual ipsi-cluster centers. Black: Mean. Red: Sigmoid fit. **d,** As **c**, but showing the mean ODI “In” ipsi-clusters for different distances at which the ODIs of neuron-pairs were swapped (local randomization control).

**Figure S4:**
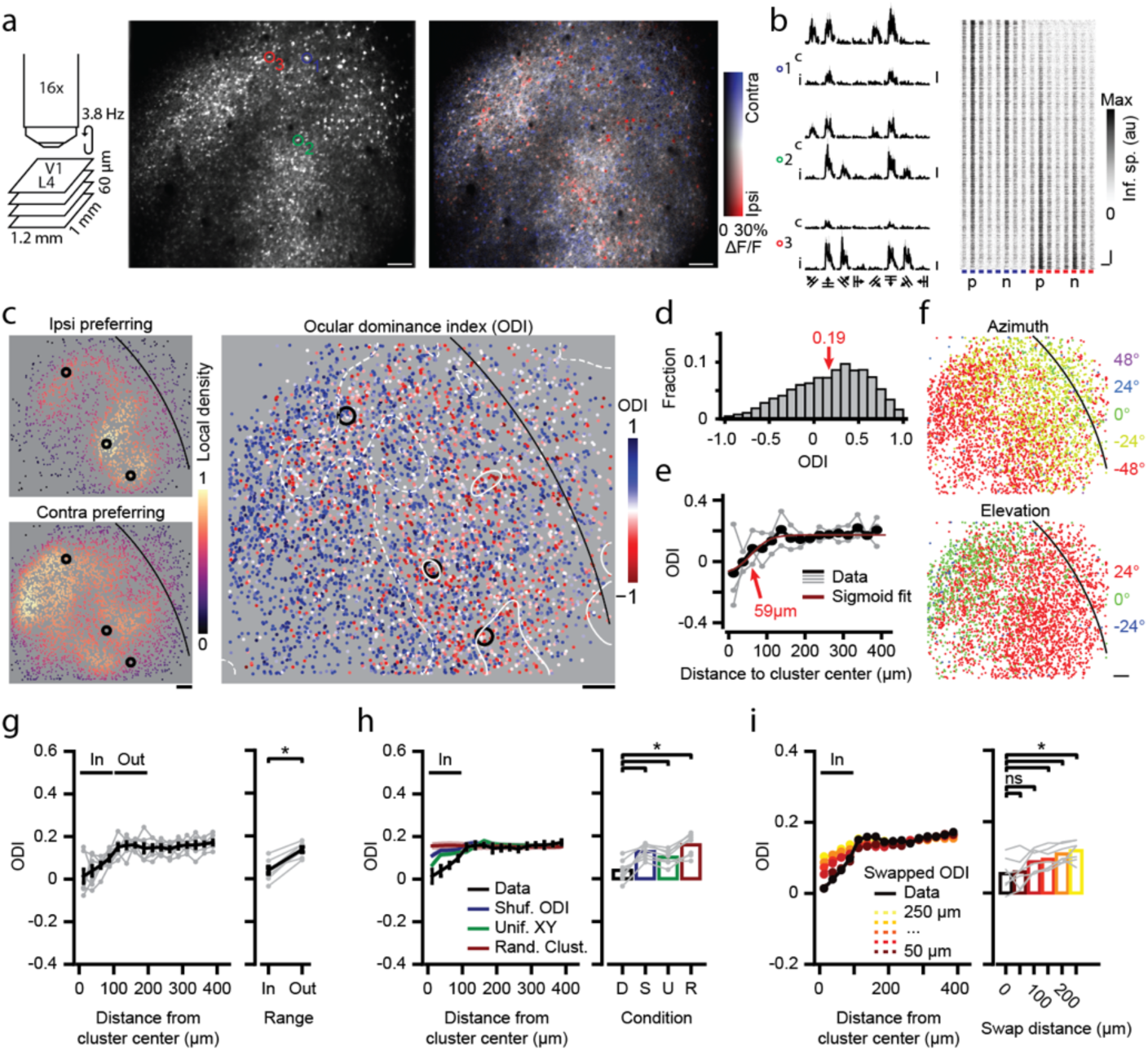
Layer 4 ipsilateral eye preferring ocular dominance clusters in Scnn1a-Tg3-Cre transgenic mice expressing the calcium indicator jRGECO1a via AAV delivery. As Fig. 1, based on imaging the red-fluorescent calcium indicator jRGECO1a in Scnn1a-Tg3-Cre mice. Using an AAV, the indicator was conditionally expressed in right-hemisphere L4 neurons. **a,** Left: Schematic of multilevel acquisition and field of view dimensions. Middle: Projection of an example motion-corrected imaging stack (single plane of a four-plane volume) showing neurons expressing jRGECO1a. Right: HLS map showing eye selective responses during visual stimulation (Hue: Preferred eye. Lightness: ΔF/F response amplitude. Saturation: Eye selectivity. Scale bar: 100 μm. **b,** Left: Trial-averaged peri-stimulus ΔF/F responses to eight directions and two eyes, for three example neurons (colored circles in **a**). Right: Trial-averaged inferred spiking responses of all visually responsive neurons (n=1794) in the imaging plane shown in **a**. **c,** Left: Local density of ipsilateral (top) and contralateral (bottom) eye preferring visually responsive neurons across the (four-plane) volume of the example mouse (M10). Black circles indicate detected ipsi-cluster centers. The black line marks the higher area boundary between V1 (left) and lateral higher visual areas (right). Right: ODI of all visually responsive neurons. White lines indicate ODI=0 (solid) and ODI=0.2 (dashed). Scale bar: 100 μm. **d,** Histogram showing the distribution of ODI values across the example volume. **e,** ODI as function of distance to detected ipsi-cluster centers for the example volume (Gray: individual centers. Black: Mean. Red: Sigmoid fit. Arrow: Point of maximum inclination, approximates the ipsi-cluster radius). **f,** Preferred azimuth (top) and elevation (bottom) of all visually responsive neurons in the example volume. Black line separates V1 (left) from lateral higher areas (right). Scale bar: 100 μm. **g,** Left: Mean ODI as function of distance to ipsi-cluster centers (Gray: individual mice). Right: ODI within a 100 μm range (“In”) of ipsi-cluster centers, compared to outside that range (“Out”, 100 μm-200 μm; two-sided WMPSR test, W=0, p=0.016, n=7 mice). **h,** Left: As **g**, but comparing original data with global randomization controls, shuffling ODI values across neurons (blue), assigning neurons new XY positions randomly sampled from a uniform distribution (green) and assigning new ipsi-cluster centers by random sampling (red). Right: Quantification, showing mean within ipsi-cluster ODI for each condition (two-sided Kruskal-Wallis test, H3=16.6, p=0.009, post hoc two-sided WMPSR test, * p<0.05, n=7 mice). **i,** As **g**, but for local randomization control, swapping the ODIs of pairs of neurons spaced apart 50 μm (dark red) to 250 μm (yellow; two-sided Kruskal-Wallis test, H5=14.0, p=0.016, post hoc WMPSR test, ns: not significant, * p<0.05, n=7 mice).

**Figure S5:**
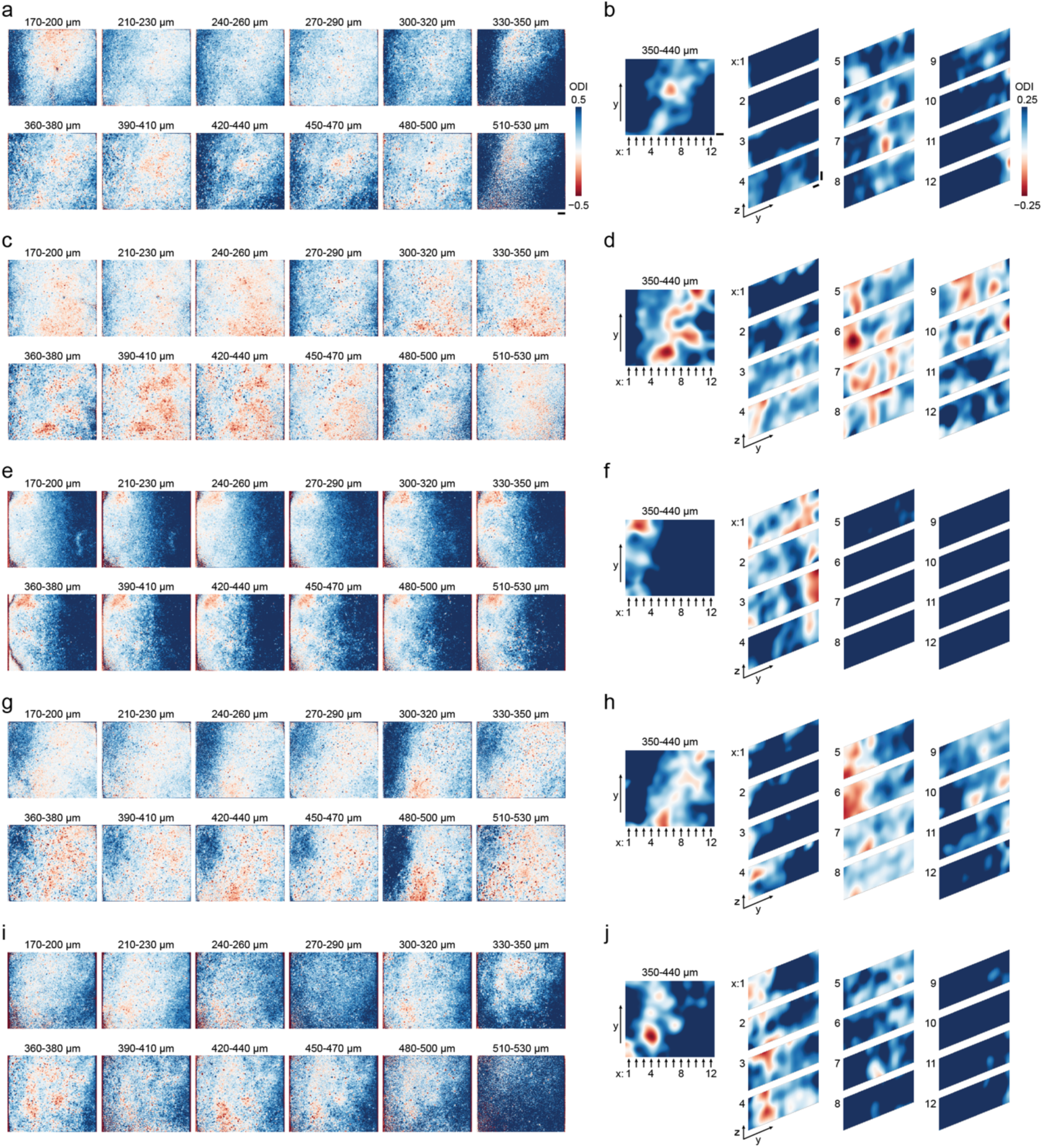
Examples of ocular dominance patterns spanning cortical layers. **a,** Pixelwise ODI maps calculated on small sub-volumes of imaging data, acquired in multiple consecutive four-plane volumes across a total depth range of 360 μm (170 μm-530 μm below cortical surface; see Fig. 2a for schematic). Scale bar: 100 μm. **b,** Left: Smoothed pixelwise ODI map of cortical layer 4 (data from 350 μm- 440 μm below cortical surface). Arrows indicate the direction along which the three-dimensional imaged volume is re-sliced to generate the side-view, smoothed ODI maps on the right. Right: Vertical slices showing smoothed pixel ODI maps spanning layer 2/3 to upper L5. Scale bar: 100 μm. **a-j,** each row represents data from a single example mouse: **a-b**, M01. **c-d**, M02. **e-f**, M03, **g-h**, M05, **i-j**, M06.

**Figure S6:**
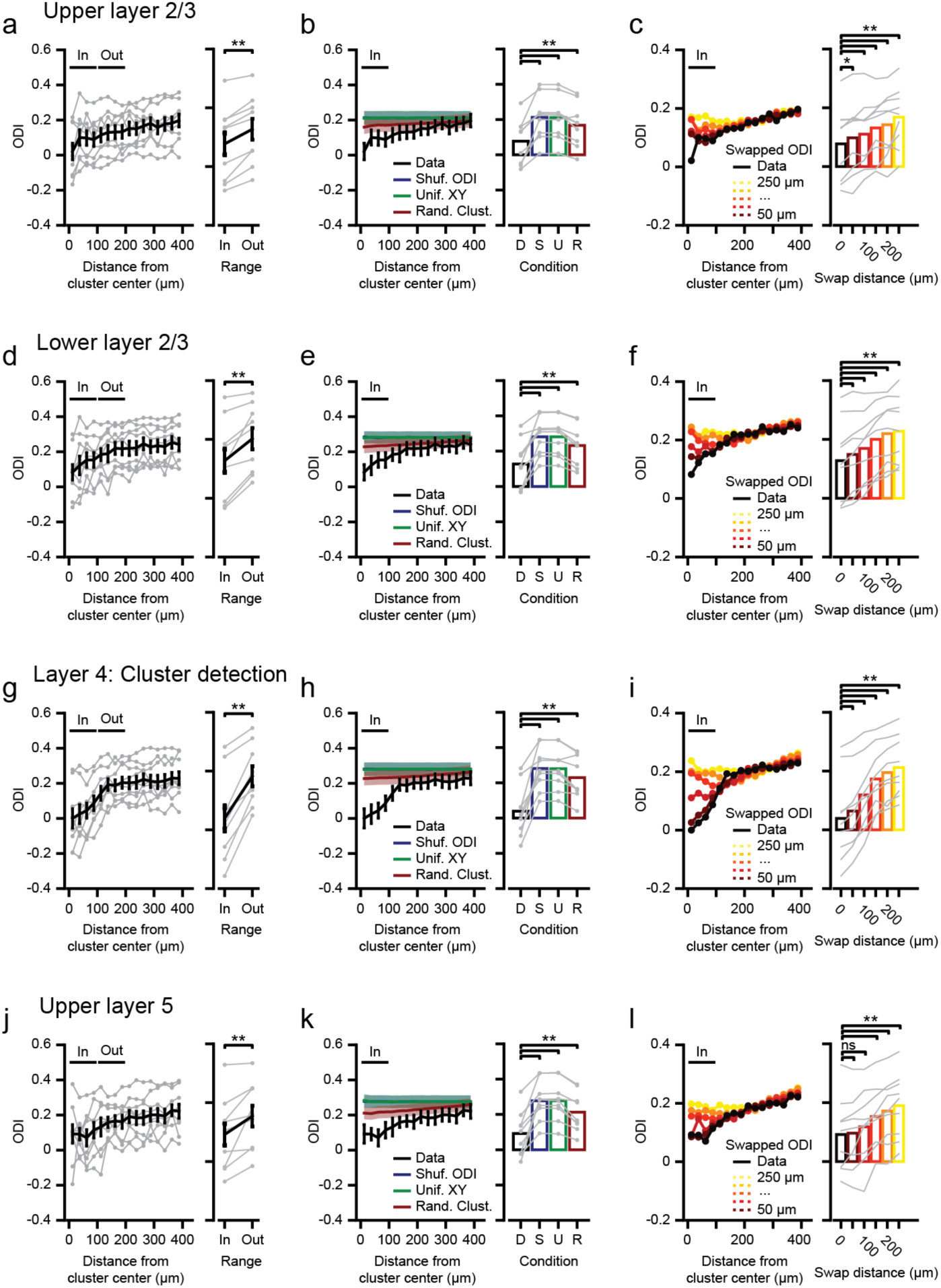
ODI of neurons in upper and lower cortical layers as a function of distance to ipsi-cluster centers identified in layer 4. **a,** Left: Mean ODI of upper L2/3 neurons as function of distance to L4-identified ipsi-cluster centers. Right: ODI within 100 μm range of L4-identified ipsi-cluster centers (“In”), compared to ODI of L2/3 neurons outside that range (“Out”; 100 μm-200 μm range). Gray: Individual mice. **b,** As **a**, but comparing ODI “In” ipsi-clusters for original data (black) to global randomization controls (blue: Shuffled ODI; green: Neuronal XY coordinates resampled from uniform distribution; red: Ipsi-cluster centers randomly sampled. **c,** As **a**, but for local randomization control, swapping the ODIs of pairs of neurons at a distance of 50 μm (dark-red) to 250 μm (yellow). **a-c**, Testing upper layer 2/3 ODI “In” ipsi-cluster centers against all controls, i.e. “Out”, global and local randomization, normalized per mouse (two-sided Kruskal-Wallis test, H9=57.2, p=4.5·10^-8^, post hoc two- sided WMPSR test, *p<0.05, **p<0.01, n=9 mice). **d-f**, as **a-c**, but for lower layer 2/3 (two-sided Kruskal-Wallis test, H9=47.9, p=2.7·10^-6^, post hoc two-sided WMPSR test, **p<0.01, n=9 mice). **g-i**, as **a-c**, but for layer 4 (two-sided Kruskal-Wallis test, H9=55.9, p=8.3·10^-8^, post hoc two-sided WMPSR test, **p<0.01, n=9 mice). **j-l**, as **a-c**, but for upper layer 5 (two-sided Kruskal-Wallis test, H9=56.3, p=7.0·10^-8^, post hoc two-sided WMPSR test, ns: not significant, **p<0.01, n=9 mice).

**Figure S7:**
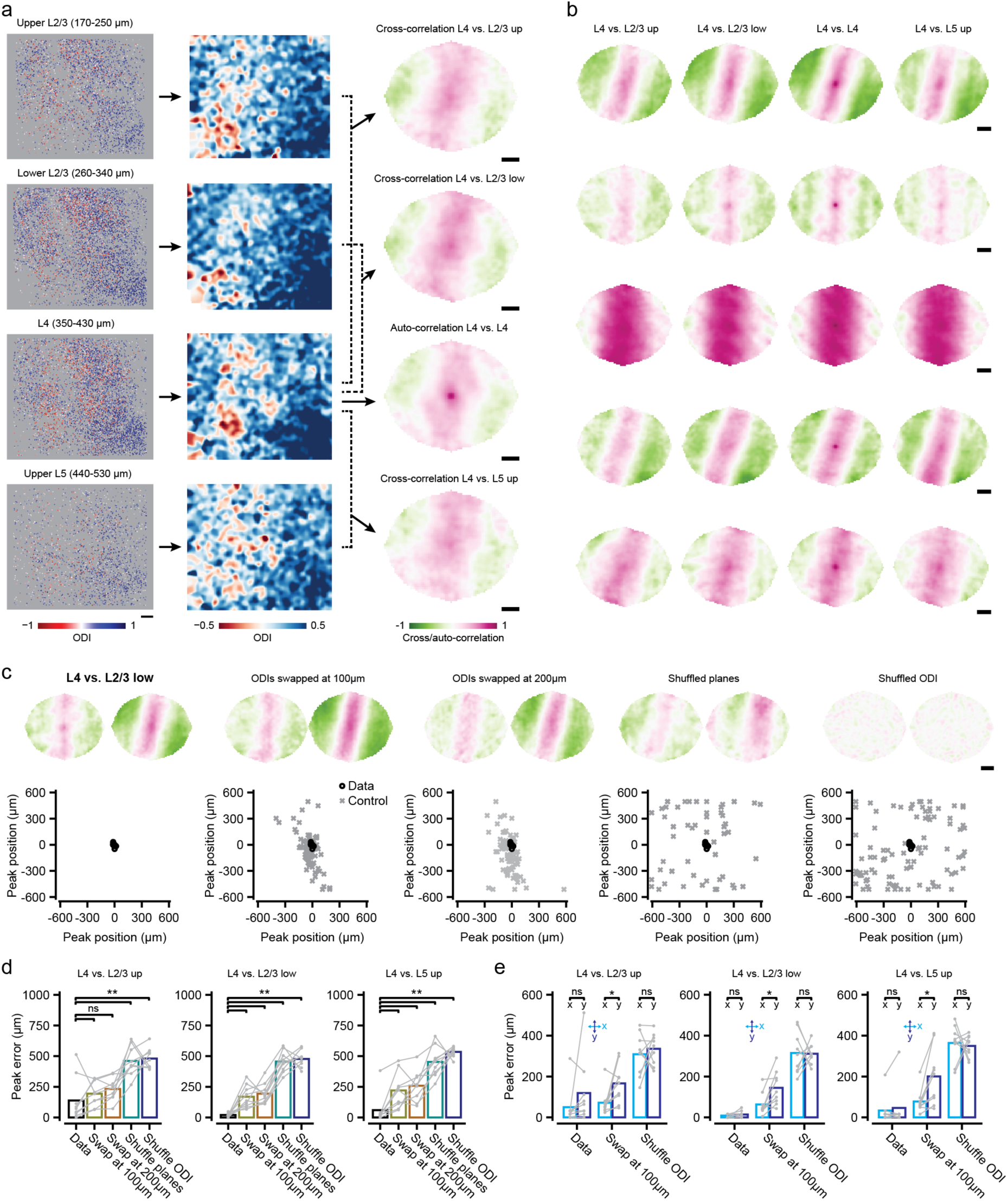
Cross-correlation analysis of ODI patterns across cortical layers reveals a local and global organization for ocular dominance. **a,** Left: For one example mouse (M06), the ODI of all visually responsive neurons in each of four depth-ranges spanning the sampled cortical volume from L2/3 to L5 (see Fig. 2a). Middle: Smoothed ODI maps (see Methods). Right: Auto-/cross-correlation of each ODI map with the L4 ODI map. **b,** Each row shows the four auto-/cross-correlation maps (as in **a**) of five further example mice (from top to bottom: M01, M02, M03, M05, M04). **c,** Left column, original data. Remaining columns: Local and global randomization controls (ODIs swapped at 100 μm, 200 μm: ODIs of random pairs of cells at approximately a distance of 100 μm or 200 μm were swapped; shuffled planes: Cross-correlation maps calculated between ODI maps of different animals; shuffled ODI: Cross-correlation maps calculated using ODI maps from data with shuffled ODI values; 10 shuffled datasets per mouse). For each column, top: Cross-correlation maps from two example mice (M02 and M01). Bottom: XY coordinates of all map peaks (original data, n=9 mice, black; shuffled data, n=9 mice, 10 shuffles per mouse, gray). **a-c,** Scale bar: 200 μm. **d,** Peak error, defined as the Euclidian distance of the cross- correlation map peak to the center of the map, for original data and local and global randomization controls (see **c**; L2/3 up: two-sided Kruskal-Wallis test, H4=27.8, p=1.4·10^-5^; L2/3 low: two-sided Kruskal-Wallis test, H4=37.8, p=1.2·10^-7^; L5 up: two-sided Kruskal-Wallis test, H4=32.0, p=1.9·10^-6^; post hoc two-sided WMPSR tests, ns: not significant, **p<0.01, n=9 mice). **e,** As **d**, but for the peak error along the horizontal and vertical dimensions of the cross-correlation maps separately (L2/3 up: two-sided Kruskal-Wallis test, H4=32.1, p=5.7·10^-6^; L2/3 low: two-sided Kruskal-Wallis test, H4=45.5, p=1.1·10^-8^; L5 up: two-sided Kruskal-Wallis test, H4=38.9, p=2.5·10^-7^; post hoc two-sided WMPSR tests, ns: not significant, *p<0.05, n=9 mice).

